# Conserved chromatin regulators control the transcriptional immune response to intracellular pathogens in *Caenorhabditis elegans*

**DOI:** 10.1101/2024.10.03.616425

**Authors:** Eillen Tecle, Samuel Li, Michael J. Blanchard, Theresa Bui, Crystal B. Chhan, Ryan S. Underwood, Malina A. Bakowski, Emily R. Troemel, Vladimir Lažetić

## Abstract

Robust transcriptional responses are critical for defense against infection. However, unrestrained immune responses can cause negative impacts such as damaging inflammation and slowed development. Here we find that a class of transcriptional regulators previously associated with regulation of development in *Caenorhabditis elegans*, is also involved in immune responses. Specifically, through forward genetics, we find that loss of *lin-15B* leads to constitutive expression of Intracellular Pathogen Response (IPR) genes. *lin-15B* encodes a transcriptional repressor with a conserved THAP domain that is associated with the DRM chromatin remodeling complex that regulates *C. elegans* development. We show that *lin-15B* mutants have increased resistance to natural intracellular pathogens, and the induction of IPR genes in *lin-15B* mutants relies on the MES-4 histone methyltransferase. We extend our analyses to other DRM and NuRD chromatin remodeling factors, as well as SUMOylation histone modifiers, showing that a broad range of chromatin-related factors can repress IPR gene expression. Altogether these findings suggest that conserved chromatin regulators may facilitate development in part by repressing damaging immune responses against intracellular pathogens.

**AUTHOR SUMMARY:** In this study, we show that transcriptional regulators, previously linked to development in *C. elegans*, also control immune responses. Through forward genetic screens, we found that loss of LIN-15B leads to constitutive activation of Intracellular Pathogen Response (IPR) genes. LIN-15B is part of the DREAM chromatin remodeling complex, and its loss enhances resistance to intracellular pathogens. This immune response depends on the MES-4 histone methyltransferase. We also discovered that other chromatin regulators, including NuRD and SUMOylation factors, similarly repress IPR gene expression, highlighting a new role in immunity for these conserved regulators of development.

## INTRODUCTION

Innate immune responses are induced by host pattern recognition receptors (PRRs) that detect pathogen-associated molecular patterns (PAMPs), leading to downstream transcriptional activation (1). One such PAMP is double-stranded RNA in the cytosol generated by RNA virus replication, which is sensed by mammalian cells using PRR RIG-I-like receptors. RIG-I-like receptors are cytosolic sensors that activate transcriptional induction of type-I interferons (IFN-I) ligands, including IFN-α and IFN-β (2–5). These secreted IFN-I ligands bind to and activate IFN-I receptors on neighboring cells, leading to a systemic IFN-I immune response (5). This response is crucial for defense against intracellular pathogens, but inappropriate IFN-I response activation can lead to developmental defects and damage due to inflammation (6–13).

Regulation of transcriptional responses involves not only transcription factors that recruit RNA polymerase to promoters, but also chromatin regulators that allow transcription factors to access regulatory regions of genes to promote transcription (14, 15). For example, the transcription factor IRF3 functions downstream of RIG-I-like receptors to induce early IFN-I ligand transcription (4), acting together with the SWI/SNF (Switch/Sucrose Non-Fermentable) complex, which is a chromatin remodeling complex that facilitates the binding of IRF3 to DNA to activate IFN-I ligand expression (16). SWI/SNF was originally identified in yeast, and is just one of many conserved chromatin remodeling complexes that regulate transcription (17, 18). There is much to be learned about how such conserved chromatin regulators control immune gene expression. and what the downstream impacts are from perturbing these regulators, particularly in a whole-animal context.

Recently, there have been similarities described in the IFN-I response, and a transcriptional immune response in the nematode *C. elegans* called the Intracellular Pathogen Response (IPR) (7). The IPR is induced by two types of intracellular pathogens that infect the *C. elegans* intestine in the wild: 1) species in the *Nematocida* genus of the Microsporidia phylum, which are obligate intracellular fungi, and 2) the Orsay virus, which is a single-stranded, positive-sense RNA virus (19–22). *C. elegans* detects viral RNA replication products from the Orsay virus using a PRR called DRH-1, which, similar to other RIG-I-like receptors, uses tandem caspase activation and recruitment domains (2CARD) to induce a downstream transcriptional program (7, 23–25). The IPR promotes resistance to both microsporidia and Orsay virus infections, and comprises an immune response distinct from those mounted against extracellular bacterial infections. Thus, similar to the anti-viral IFN-I response in mammals, the IPR is a transcriptional response in *C. elegans* that provides defense against intracellular pathogens.

While the IPR promotes protection against intracellular pathogens, its activation can come at the cost of impaired organismal health in the absence of infection, similar to overactivation of the IFN-I response (7, 21, 22, 26). For example, mutations in the *C. elegans* gene *pals-22* (from the *pals* gene family of unknown biochemical function) lead to constitutive IPR gene expression as well as increased resistance to infection by intracellular pathogens (21, 22). However, *pals-22* mutants have slowed growth, shortened lifespan, and signs of neuronal dysfunction (21, 22). These phenotypes can be reversed by a mutation in the downstream positive regulator *pals-25*, which activates IPR gene expression in a *pals-22* mutant background (22, 27). This *pals-22/pals-25* module appears to act as an OFF/ON switch for IPR gene expression separately from DRH-1 activation of the IPR, and separately from another *pals* gene module comprised of *pals-17/pals-20/pals-16* identified through forward genetic screens for regulation of IPR transcription (22, 26–28). Similar to *pals-22* mutants, *pals-17* mutants have constitutive IPR gene expression and increased resistance against intracellular pathogens, but impaired development (26).

While several genes in *C. elegans* like *pals-22/25* and *pals-17/16/20* have been shown to regulate development, IPR gene expression and resistance to infection (22, 26, 27, 29), we know relatively little about downstream factors that mediate transcriptional induction, such as the direct regulators of IPR gene transcription. For example, while DRH-1 is a homolog of RIG-I-like receptors, *C. elegans* lacks homologs for several factors acting downstream of these receptors, such as the transcription factor IRF3. Instead, an RNAi screen in *C. elegans* identified the bZIP transcription factor ZIP-1 to be acting downstream of DRH-1, as well as downstream of other IPR triggers like *N. parisii* infection and *pals-22* mutations (30). ZIP-1 promotes the early expression of *pals-5*, which is commonly used as a reporter gene for IPR induction. Altogether ZIP-1 is responsible for activating about 1/3 of IPR genes, and promotes resistance to intracellular pathogens (30). However, the transcription factor responsible for inducing *zip-1*-independent IPR genes is unknown, and there is much to be learned about how transcriptional regulators such as chromatin remodeling complexes and histone modifiers may affect IPR transcription.

Here we perform two independent forward genetic screens that both converge on the chromatin regulator LIN-15B as a new negative regulator of the IPR. LIN-15B is a THAP-domain (Thanatos-associated protein domain) protein and is functionally associated with the chromatin remodeling DREAM complex (DRM complex in *C. elegans*), which plays a key role in transcriptional repression during development. Our findings reveal that *lin-15B* functions upstream of the gene *mes-4*, which encodes an H3K36 methyltransferase histone modifier crucial for chromatin remodeling (31). Importantly, we show that, together with its activation of the IPR, loss of LIN-15B provides enhanced protection against infection with both the Orsay virus and *Nematocida parisii* microsporidia. LIN-15B, along with DRM complex genes, is part of the broader SynMuv (synthetic multivulva) family of developmental genes, which also includes Nucleosome Remodeling and Deacetylase complex (NuRD) genes and genes involved in SUMOylation. By testing various members of the SynMuv family, we found that most SynMuv genes, including both chromatin remodelers and histone modifiers, negatively regulate the IPR. In summary, our findings revealed the involvement of different conserved chromatin remodeling and transcription-regulating complexes in modulating immunity against obligate intracellular pathogens, and raise the intriguing possibility that several of these factors may allow for normal development by repressing expression of innate immune genes.

## RESULTS

### Two independent genetic screens isolate DRM-associated protein LIN-15B as a novel negative regulator of the IPR

To uncover novel negative regulators of the IPR, we conducted two independent forward EMS mutagenesis screens using the *pals-5p::gfp* transcriptional IPR reporter. In both screens, we identified mutants with constitutive expression of *pals-5p::gfp* in the absence of infection. The first screen (33,504 haploid genomes screened) was performed in the *pals-22 pals-25* mutant background, which has wild-type levels of IPR expression and induction, due to loss of both the IPR negative regulator *pals-22* and the IPR activator *pals-25*. The screen was performed in this genetic background in order to not identify new mutations of *pals-22*. Four mutants with constitutive *pals-5*p::GFP expression were isolated in this screen, one of which was the IPR negative regulator *pals-17*, as previously published (26). The second screen was performed in a wild-type genetic background (26,000 haploid genomes screened) and nine mutants with constitutive *pals-5*p::GFP expression were isolated in this screen, including the IPR negative regulator *pnp-1*, as previously published (29).

After performing complementation analysis among mutants from these two screens, we found that one mutant from each of the two screens failed to complement each other for the phenotype of constitutive *pals-5*p::GFP expression, suggesting that they may encode mutations in the same gene. Whole genome sequencing analyses revealed that both of these mutants had mutations in the gene *lin-15B*. The mutant from the first screen contained three missense mutations in the second exon of *lin-15B*, leading to the following amino acid changes: Ala56Thr, His157Gln, and Ala168Thr, and was designated the *jy76* allele of *lin-15B*. The mutant from the second screen had a single missense mutation in the seventh exon of *lin-15B*, resulting in a Cys1142Tyr substitution in the protein sequence, and was designated the *jy86* allele of *lin-15B* (Fig 1). This mutation is located in the region that encodes a THAP domain, protein motif with similarity to the DNA-binding domain of P element transposase (32), which is important for LIN-15B function as a transcriptional regulator (33).

**Fig 1.**
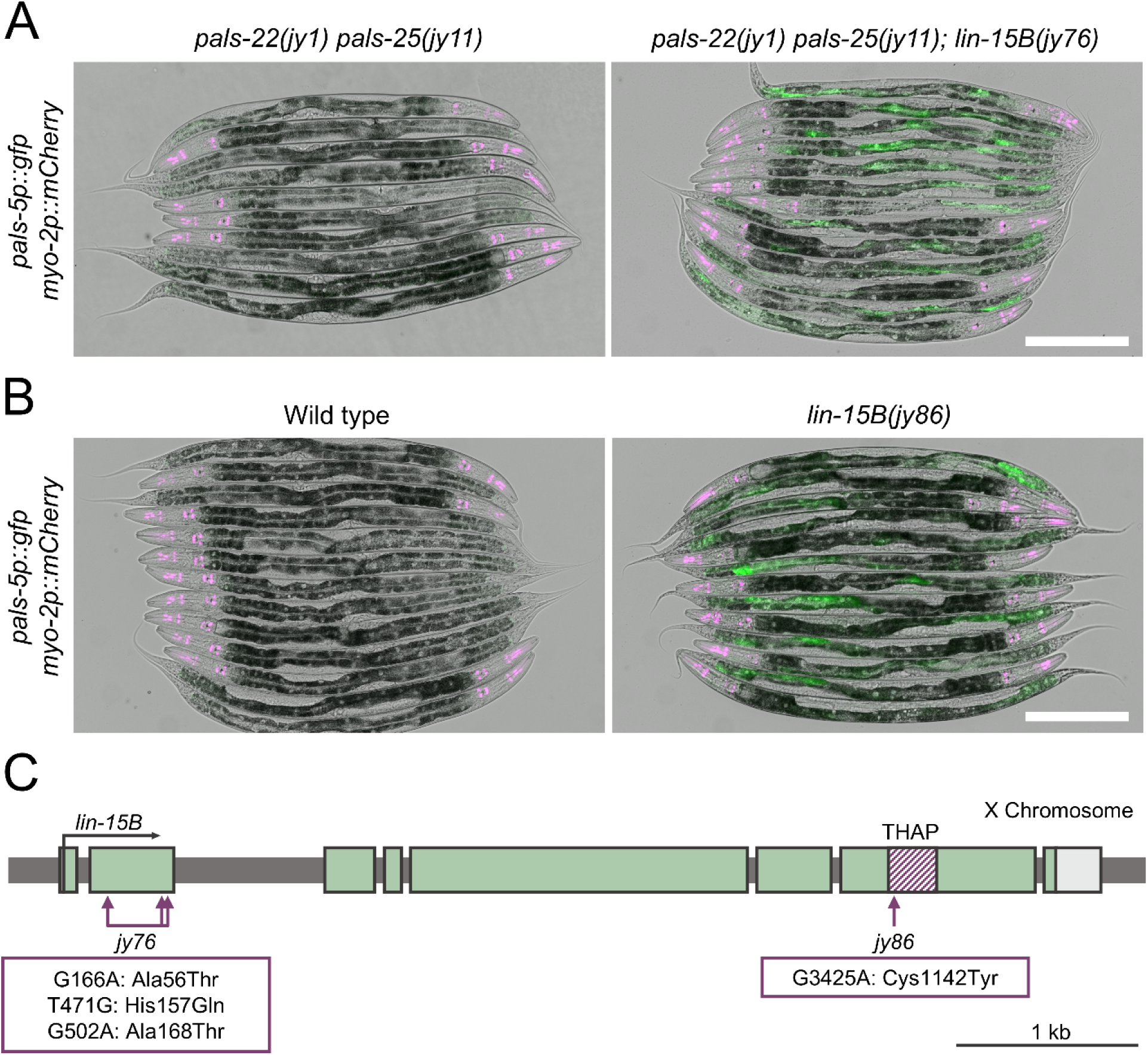
*lin-15B* is a negative regulator of *pals-5*p::GFP expression. (A, B) Mutants isolated from two independent forward genetic screens in *pals-22 pals-25* double mutant (A) and wild-type backgrounds (B) show increased expression of *pals-5*p::GFP reporter (shown in green). *myo-2*p::mCherry co-injection marker is shown in magenta. Green, magenta and DIC channels were merged. Scale bar = 200 µm. (C) *lin-15B* gene structure. *lin-15B* exons are indicated with green boxes, UTRs are shown in light gray, region encoding THAP domain is indicated with striped magenta box. Horizontal arrow shows the direction of transcription. Vertical arrows indicate positions of point mutations in alleles *jy76* and *jy86*.

To determine whether endogenous *pals-5* mRNA levels are increased in *lin-15B* mutants and to identify the tissues in which it is expressed, we performed single-molecule fluorescence in situ hybridization (smFISH) analysis in *lin-15B(jy76)* mutants. While *pals-5* mRNA was undetectable in wild-type animals, *lin-15B* mutants exhibited numerous *pals-5* mRNA puncta (Fig 2A). Additionally, *pals-5* mRNA expression appeared to be specific to intestinal cells in this mutant, even though *pals-5* can be expressed more widely in other genetic backgrounds, such as when *pals-22/25* signaling is perturbed. However, only a subset of intestinal cells expressed *pals-5* puncta in *lin-15B* mutants, and the pattern of expression seemed to be stochastic, as different intestinal cells showed *pals-5* mRNA expression in different animals. In summary, *lin-15B* mutants had not only *pals-5* reporter induction but also endogenous *pals-5* mRNA induction.

**Fig 2.**
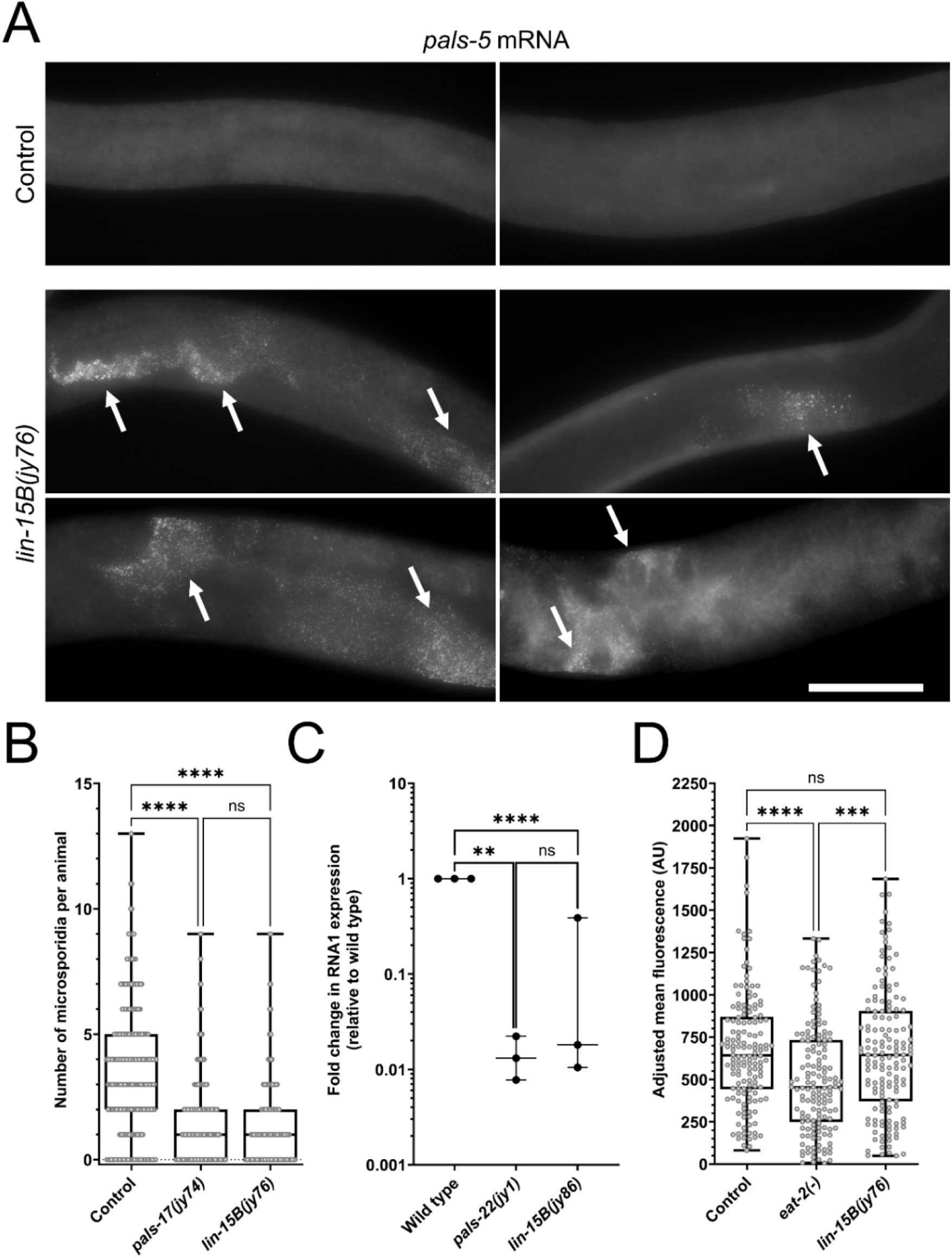
IPR activation in *lin-15B* mutants provides protection against natural intracellular pathogens of the intestine. (A) *pals-5* mRNA expression is induced in intestinal cells in *lin-15B* mutants relative to control strain. Z-stack maximal projections are shown. Arrows indicate regions with *pals-5* mRNA puncta. Scale bar = 50 µm. (B) *N. parisii* pathogen load quantified at 3 hpi as number of sporoplasms per animal. (C) qRT-PCR analysis of Orsay virus pathogen load. Each experimental replicate is shown by a dot. A one-tailed t-test was used to calculate *p*- values; asterisks represent a significant difference between the labeled sample and the wild-type control; **** *p* < 0.0001; ** *p* < 0.01; ns = no significant difference. (D) Box-and-whisker plot of adjusted bead fluorescence levels per animal. The *eat-2(ad465)* mutant strain is a feeding-defective control. AU = arbitrary units. (B, D) Box lines represent median values, while box edges mark the 25^th^ and 75^th^ percentiles. Whiskers extend to the minimum and maximum values. Three experimental replicates were performed. Gray circles indicate individual values for each animal; 100 (B) and 50 (D) animals were analyzed per strain per replicate. The Kruskal-Wallis test was used to determine *p*-values: **** *p* < 0.0001; *** *p* < 0.001; ns = no significant difference.

### *lin-15B* mutants have increased immunity against natural intracellular pathogens of the intestine

Because IPR induction has previously been linked to increased resistance to intracellular pathogens (22, 26, 29, 30), we tested whether *lin-15B* mutants exhibit enhanced immunity against these pathogens. First, we examined the response to *N. parisii*, which is the most commonly found microsporidia species infecting *C. elegans* in the wild, and has tropism for the intestine (28, 34). Using a well-established feeding method, we infected the strain in which *jy76* allele was isolated (negative control) *pals-17* mutants from the same screen (positive control), and *lin-15B(jy76)* mutants with microsporidia *N. parisii* spores and quantified the pathogen load at the sporoplasm (early parasite cell) stage across these three backgrounds. Here we found that *lin-15B* mutants had a lower number of sporoplasms in the intestine per animal, compared to the negative control, showing similar resistance to the *pals-17* mutants. Thus, *lin-15B* appears to regulate immunity against *N. parisii* infection (Fig 2B).

Next, we investigated whether *lin-15B* mutants had increased resistance to infection with the Orsay virus, which is also a natural pathogen of *C. elegans* with tropism for the intestine. Using a feeding method to introduce pathogen infection, we infected wild-type animals (with *pals-5p::gfp* reporter; negative control), *pals-22* mutants (positive control), and *lin-15B(jy86)* mutants with the Orsay virus and measured Orsay RNA1 levels using qRT-PCR. Here our results showed a significant decrease in viral RNA in *lin-15B* mutants, comparable to the reduction seen in *pals-22* mutants, again supporting the model that *lin-15B* regulates immune responses against intracellular pathogens (Fig 2C).

To rule out the possibility that differences in food intake or excretion might explain the variations in pathogen load in *lin-15B* mutants, we measured intestinal accumulation of fluorescent beads in the original strain used in the screen, *eat-2*(*-*) eating-defective control and *lin-15B* mutants. Here we found that *lin-15B* mutants accumulated beads similarly to the original strain, suggesting that the enhanced immunity in *lin-15B* mutants is not due to altered feeding behavior (Fig 2D). In conclusion, our data indicate that the loss of *lin-15B* promotes immunity against natural intracellular pathogens of the *C. elegans* intestine.

### Loss of LIN-15B induces multiple IPR genes, dependent on histone methyltransferase MES-4

To determine the extent of IPR gene expression regulated by *lin-15B*, we analyzed two previously published transcriptomic data sets for *lin-15B* loss-of-function mutant animals (33, 35). Here we found that 69 out of 80 canonical IPR genes (86.25%) were induced in at least one analyzed data set, with 42 IPR genes (52.5%) being induced in both data sets (including *pals-5*) (Fig 3A, Table S1). Similar to previously identified mutants with constitutive upregulation of the IPR, like *pals-22* and *pals-17* mutants (21, 26), both *lin-15B* mutant alleles we isolated displayed developmental delay. We found that a synchronized population of *lin-15B* mutants takes 10 hours longer than control strains to progress from the first larval stage (L1) to the fourth larval stage (L4) when grown at 20°C (58 h vs. 48 h). Due to the presence of a THAP domain in LIN-15B, which is a DNA-binding motif involved in regulating gene transcription and chromatin structure (36, 37), we hypothesized that this protein might directly bind to IPR genes whose expression it regulates. However, previous ChIP-seq studies of LIN-15B (33) revealed that it only directly binds to five IPR genes (*pals-26*, *pals-27*, *Y6E2A.5*, *fbxa-75* and *Y41C4A.11*) (Table S2), suggesting that it might indirectly regulate transcription of the majority of IPR genes.

**Fig 3.**
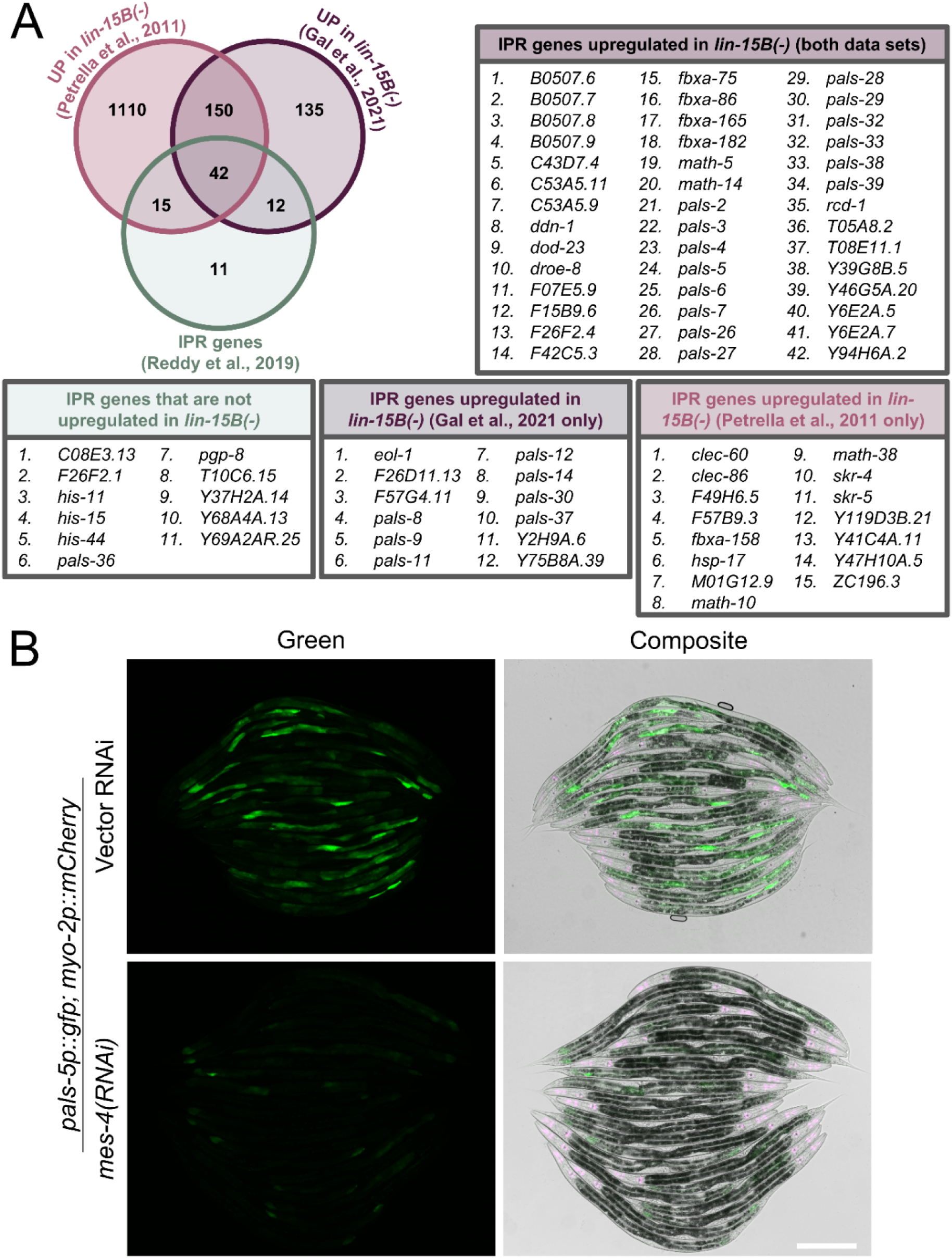
The majority of IPR genes are upregulated in *lin-15B* mutants and *pals-5*p::GFP expression is dependent on *mes-4*. (A) *In silico* analysis of two previously published transcriptomic datasets for *lin-15B* mutants. (B) *pals-5*p::GFP expression in *lin-15B* mutants is reduced following *mes-4* RNAi treatment. *pals-5*p::GFP is shown in green; *myo-2*p::mCherry co-injection marker is shown in magenta. Magenta, green and DIC channels were merged in the composite images. Scale bar = 200 µm.

LIN-15B is a protein associated with the dimerization partner, RB-like, E2F and multi-vulval class B (DREAM) complex, called DRM in *C. elegans* (33). Thus, we refer to LIN-15B as a DRM-AP for DRM-associated protein. The retinoblastoma (Rb) ortholog in *C. elegans* is named LIN-35, and is associated with development (33, 38–43). Our analysis of published transcriptomic data from *lin-35* mutants (35) revealed that the majority of IPR genes (62.5%; 50/80) are functionally repressed by LIN-35 (S1 Fig). LIN-35 is a transcriptional repressor that among other functions, prevents expression of histone methyltransferase MES-4 in somatic tissues including the intestine (35, 44, 45). Interestingly, MES-4 is important for expression of many genes that are repressed by LIN-35 (35). We analyzed if *lin-35*-dependent IPR genes are *mes-4*-dependent and found that 70% (35/50) of those genes or 43.75% (35/80) of all IPR genes require *mes-4* for their transcriptional upregulation in *lin-35* mutant background (S1 Fig). Therefore, our analysis suggests that *lin-35* predominantly suppresses the induction of IPR genes by downregulating or inhibiting *mes-4* expression.

Next, we investigated whether *lin-15B* requires *mes-4* for IPR induction. To address this question, we analyzed *pals-5*p::GFP expression in *lin-15B(jy76)* mutants on control and *mes-4* RNAi plates. We observed a substantial reduction in GFP expression following *mes-4* RNAi treatment, suggesting that, similar to *lin-35* mutants, IPR induction in *lin-15B* mutants is *mes-4*-dependent (Fig 3B). Previous ChIP-seq studies revealed that *mes-4* is one of the genes bound by LIN-15B (S2 Table) (33), potentially indicating a mechanism for IPR gene regulation.

Although *mes-4* mRNA expression was not upregulated in *lin-15B* mutants from the same study (33), an earlier transcriptomic study of *lin-15B* mutants reported an increase in *mes-4* mRNA levels (S1 and S2 Tables) (35). By analyzing the same ChIP-seq dataset (33), we found that LIN-15B binds to *pals-17* and *pals-22*, two negative regulators of the IPR (S2 Table). However, mRNA expression of these genes was not significantly altered in *lin-15B* mutants (S1 Table) (33, 35), perhaps due to low expression level of these two genes. In conclusion, we found that LIN-15B regulates IPR gene expression, possibly via suppression of *mes-4*.

### Loss of multiple chromatin regulators (SynMuv genes) induce the IPR induction

*lin-15B* and *lin-35* are members of the SynMuv family of genes that regulate vulval development as well as other developmental processes through their involvement in transcriptional regulation and chromatin regulation (Fig 4A) (46–48). Using an RNAi approach, we investigated whether the downregulation of other SynMuv genes also induces expression of the IPR. We analyzed *pals-5*p::GFP reporter expression following RNAi of different SynMuv genes in two different age categories – L1 to L3 stage and L4 to adult stage. We classified GFP expression as none, weak, and strong GFP. L4440 control vector and *pals-17* RNAi were used as negative and positive controls for IPR induction, respectively.

**Fig 4.**
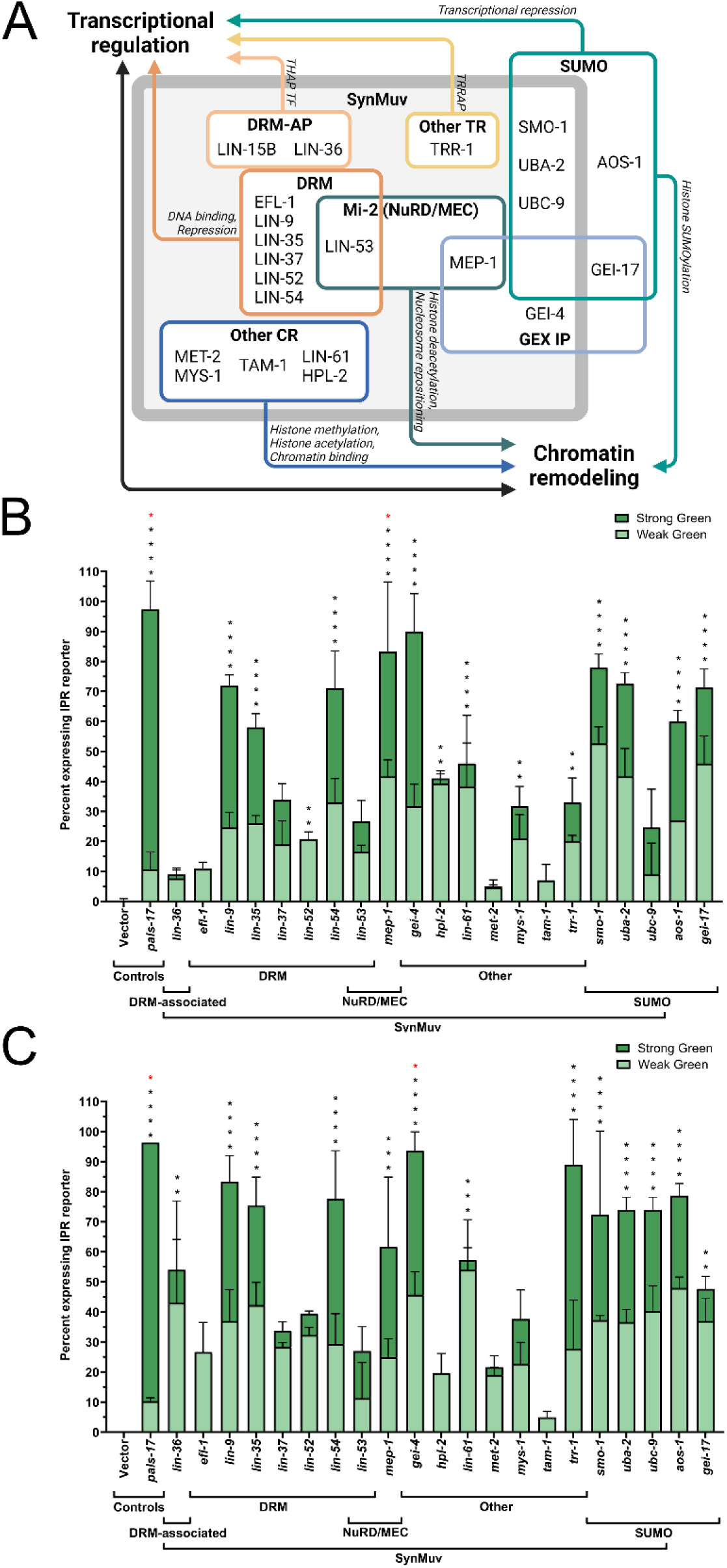
Multiple chromatin regulators in the SynMuv class negatively regulate *pals-5*p::GFP expression. (A) Model depicting functional complexes in the SynMuv gene/protein family and their roles in transcriptional regulation and chromatin remodeling. Abbreviations: DRM – DP, Rb, Myb; DRM AP – DRM-Associated Proteins; TR – Transcription Regulators; NuRD – Nucleosome Remodeling and Deacetylase; GEX – Gut on EXterior; IP – Interacting Protein; CR – Chromatin Remodeler. (B, C) Downregulation of different SynMuv components induces *pals-5*p::GFP expression in L1-L3 (B) and L4-Adult developmental stages (C). Error bars are shown separately for the “Strong GFP” and “Weak GFP” categories and represent SD. Statistical analyses were performed on the total number of green animals (both weak and strong GFP expression), with all values compared to the control vector RNAi. Black asterisks represent *p*-values calculated using ordinary one-way ANOVA test with Holm-Šídák multiple comparison test; red asterisks represent *p*-values calculated using the Kruskal-Wallis test; **** *p* < 0.0001; *** *p* < 0.001; ** *p* < 0.01; * 0.01 < *p* < 0.05; *p*-values higher than 0.05 are not labeled.

We found that downregulation of multiple SynMuv genes from different functional subfamilies induces the expression of the IPR reporter (Figs 4B and 4C). In general, stronger induction was observed in the older age category of the animals. Our results suggest that SynMuv genes encoding various components of the DRM complex and DRM-APs negatively regulate the IPR. The DRM complex is composed of several key components: a repressive transcription factor EFL-1 protein with its dimerization partner DPL-1, an Rb-like pocket protein LIN-35, and a core MuvB complex consisting of five subunits – LIN-9, LIN-37, LIN-52, LIN-54, and LIN-53 (46, 49). Interestingly, we observed varying levels of GFP expression across different DRM RNAi treatments. For example, downregulation of *lin-54*, *lin-35*, and *lin-9* caused the strongest induction of the IPR reporter, while *efl-1* downregulation resulted in a more modest effect (Figs 4B and 4C). Additionally, downregulation of *lin-36*, which encodes a DRM-AP with a THAP domain, led to low induction of *pals-5*p::GFP in early larval stages, but its expression increased in L4 and adult animals (Figs 4B and 4C).

To explore whether these differences stem from RNAi efficacy or actual functional differences between the genes, and to gain further insights into the expression changes of multiple IPR genes, we analyzed transcriptomics data from previous studies on several DRM mutants (33). We found that, in addition to *lin-15B* and *lin-35* mutants, a subset of IPR genes is upregulated upon the loss of other DRM genes, such as *dpl-1* (10%, 8/80) and *lin-37* (57.5%, 46/80) (S2 Fig and S3 Table). However, no significant upregulation of IPR gene expression was observed in *efl-1* mutants. Likewise, no IPR gene induction was present in *lin-36* mutants (S2 Fig and S3 Table). These findings align with our RNAi studies, where we observed stronger *pals-5*p::GFP expression in *lin-35(RNAi)* and *lin-37(RNAi)* animals, and lower GFP expression in *efl-1(RNAi)* and *lin-36(RNAi)* animals (Figs 4B and 4C). Taken together, these data suggest that different DRM components may have distinct roles in regulating IPR induction.

In our RNAi screen, we also observed that components of complexes containing the Mi-2 nucleosome remodeler, NuRD and MEP, negatively regulate IPR reporter expression (Fig 4). RNAi of several other SynMuv genes also robustly induced *pals-5p::GFP* expression, particularly when *gei-4* (encoding GEX Interacting protein) and *trr-1* (encoding TRRAP-like protein) were downregulated (Fig 4). However, downregulation of *met-2* (encoding histone methyltransferase) and *tam-1* (Tandem Array expression Modifier; involved in ubiquitylation) had relatively weak or almost no effect on IPR reporter expression (Fig 4). We also analyzed available transcriptomics data for *met-2* mutants and found no induction of IPR genes expression in this genetic background (S3 Table). In summary, our results suggest that a subset of SynMuv genes play a role in IPR induction.

### Loss of SUMOylation factors induces multiple IPR genes

Several genes encoding core SUMOylation components are also part of the SynMuv gene family (47, 50). SUMOylation is a conserved post-translational modification process that alters the function of many substrate proteins and is closely linked to transcriptional regulation and chromatin remodeling (51–54). In *C. elegans*, this process involves the stepwise attachment of a small protein, SUMO (or SMO-1), to a substrate via a cascade of enzymatic activities involving E1 (composed of AOS-1 and UBA-2), E2 (UBC-9), and E3 complexes (GEI-17) (55, 56). In our RNAi screen, we found that downregulation of all three SynMuv genes from the SUMO family – *smo-1*, *uba-2*, and *ubc-9* – induced the expression of the *pals-5*p::GFP reporter (Figs 4B, 4C and 5A). We also discovered that SUMOylation genes *aos-1* and *gei-17*, not previously annotated as SynMuv, also negatively regulate the IPR (Figs 4B and 4C).

To assess whether SUMOylation affects the endogenous expression of *pals-5* and several other IPR genes, we conducted qRT-PCR analysis. We observed a substantial increase in IPR gene expression following RNAi treatments targeting *smo-1*, *uba-2*, and *aos-1*, with modest or low IPR induction following the downregulation of other SUMOylation genes (Fig 5B). Our previous yeast two-hybrid analysis demonstrated that the positive IPR regulator PALS-25 physically interacts with three SUMOylation components involved in IPR regulation: SMO-1, UBC-9, and GEI-17 (27). To explore this connection further, we conducted a similar analysis using the IPR activator PALS-20 as bait and identified a subset of shared interactors with PALS-25. Notably, all three SUMOylation proteins were found to interact with both PALS-20 and PALS-25 (Fig 5C and S4 Table). In summary, our findings suggest a novel role for SUMOylation in regulating the transcription of IPR genes, possibly by modulating the activity of PALS proteins as positive regulators of the IPR.

**Fig 5.**
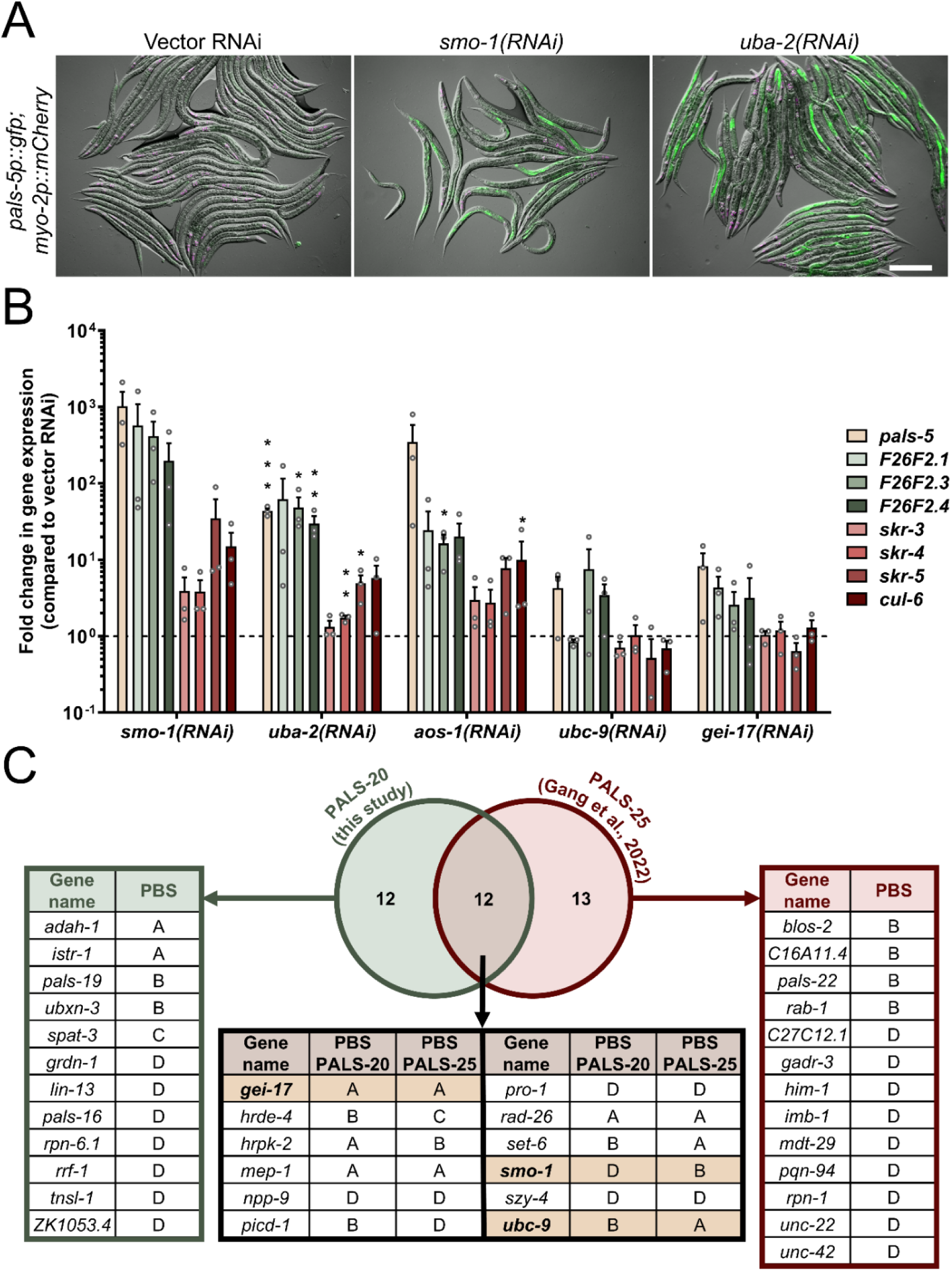
Depletion of SUMOylation components induces IPR gene expression. (A) *smo-1* and *uba-2* RNAi treatments induce *pals-5*p::GFP expression in intestine. Green, magenta and DIC channels were merged. *pals-5*p::GFP is shown in green; *myo-2*p::mCherry co-injection marker is shown in magenta. Scale bar = 200 µm. (B) qRT-PCR analysis of representative IPR genes following downregulation of major SUMOylation components. Gene expression shown as the mean fold changes relative to the wild-type control in mixed stage population. Error bars are SD. A one-tailed t-test was used to calculate *p*-values; asterisks represent a significant difference between the labeled sample and the vector control; *** *p* < 0.001; ** *p* < 0.01; * *p* < 0.05; p-values higher than 0.05 are not labeled. (C) Yeast two-hybrid analysis of PALS-20 and PALS-25 interactors, including SUMOylation genes SMO-1, UBC-9 and GEI-17 (yellow). PBS: A = very high confidence in the interaction, B = high confidence in the interaction, C = good confidence in the interaction, D = moderate confidence in the interaction

## DISCUSSION

Mechanisms of chromatin remodeling during the activation of transcriptional immune responses are not well understood. In this study, we identified several chromatin remodeling and transcriptional repressor complexes that act as negative regulators of the IPR in *C. elegans* (Fig 6). First, through an unbiased approach, we discovered that *lin-15B* functions as a repressor of the IPR. We demonstrated that the loss of *lin-15B* confers resistance to obligate intracellular pathogens. Our RNAi screen revealed that multiple genes from the SynMuv family, which are functionally related to *lin-15B*, also negatively regulate the IPR. Mechanistically, our data suggest that *lin-15B* operates upstream of *mes-4*, which is necessary for IPR induction in this context (Fig 6).

**Fig 6.**
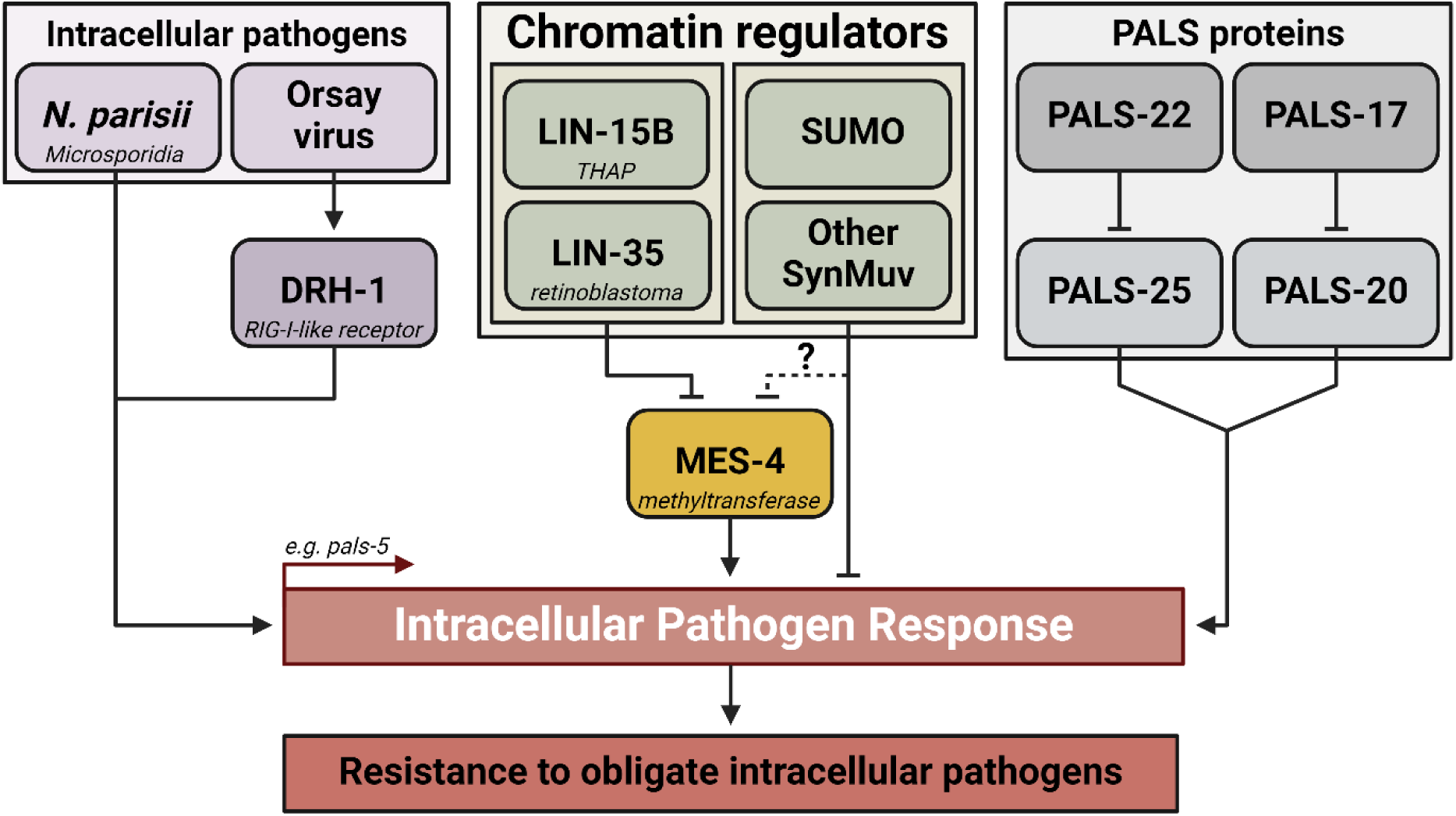
Model of IPR regulation by SynMuv genes and *mes-4*.

Although LIN-15B lacks a direct mammalian homolog (57), it features a well-conserved THAP domain, which is frequently found in transcriptional regulators (33, 58). While this domain is not essential for DNA binding by LIN-15B, it is crucial for its functionality (33). LIN-15B, along with another SynMuv THAP-domain protein, LIN-36, is functionally linked to the conserved DRM complex, which globally represses gene expression and inhibits cell cycle progression in differentiated cells (33, 59, 60). The DRM complex uses these THAP proteins to regulate its genomic targets; however, LIN-15B and LIN-36 govern different subsets of DREAM-regulated genes via distinct mechanisms (33). LIN-15B primarily represses germline genes by promoting H3K9me2 tagging at their promoters, while LIN-36 represses cell-cycle genes by enriching the histone variant H2A.Z within gene bodies (33). By analyzing expression of IPR fluorescent reporter following *lin-36* depletion, we observed some induction of the GFP especially in older animals, suggesting that roles of *lin-15B* and *lin-36* might somewhat overlap in this context. However, based on transcriptomic data, *lin-36* does not seem to function as IPR repressor (33).

As for the role of DRM complex components’ in IPR regulation, we found that some components of this complex have stronger effect on IPR regulation than others. This result could mean that not all components of DRM complex are involved in transcriptional repression of the IPR and that a subset of DRM genes might form a distinct complex involved in immune regulation. For example, we found that *efl-1* RNAi treatment had a minor effect on *pals-5*p::GFP expression induction and that none of IPR genes are induced in an *efl-1* mutant background. Interestingly, EFL-1 is essential for repressing LIN-35 targets shared with LIN-36, but it is not required for repressing those shared with LIN-15B (33). Alternatively, certain components of the DRM complex might have redundant roles, and therefore downregulating single subunit would not cause drastic effect on IPR induction.

Analysis of previously published microarray, RNA-seq, and ChIP-seq datasets revealed that LIN-15B regulates expression of most IPR genes but binds to DNA for only a few of them (33, 35). Additionally, LIN-15B binds *mes-4* gene and requires this gene for IPR reporter induction. MES-4 is a histone methyltransferase that catalyzes lysine 36 methylation on histone 3 (31, 61), an epigenetic mark characteristic of transcriptionally accessible chromatin (62, 63). These regions may contain IPR genes, suggesting that LIN-15B might regulate IPR gene expression indirectly through the repression of MES-4. *mes-4* is a single homolog of three closely related mammalian genes Nsd1, Nsd2 and Nsd3 (Nuclear receptor-binding SET domain protein) (61). MES-4 promotes transcription of germline genes and is most highly expressed in the germline (58, 64). Some DRM components (LIN-54/LIN54 and EFL-1/E2F) and MES-4 co-bind the same genomic targets in the germline, fine-tuning gene expression and allowing the proper development of the germline cells (65). Interestingly, LIN-35 shows reduced binding in the germline, whereas, in somatic tissues, it directly binds to the promoter region of *mes-4* and represses its transcription (44). Loss of *lin-35* leads to transcriptional upregulation of *mes-4* in the soma, initiating the germline fate in these tissues (35, 44, 45). Future studies will investigate if similar changes occur during infection with obligate intracellular pathogens.

Our data demonstrate that *lin-15B* mutants have increased resistance to infection with the intestinal pathogens microsporidia and the Orsay virus. This finding aligns with recent findings suggesting that the activity of SynMuvB genes is downregulated during Orsay virus infection (66), and with studies indicating that *lin-35* mutants confer maternally transferred immunity against microsporidia (67). One possibility is that these pathogens may inhibit SynMuv components like LIN-35/Rb and LIN-15B in intestinal cells to induce a less differentiated state, promoting faster replication in terminally differentiated cells, which often contain less deoxynucleotides and other precursors required by viruses and microsporidia for growth (68). Notably, a comparative analysis of microsporidian genomes revealed that all examined species encode DP and E2F components of the DREAM complex but lack Rb (69). It is intriguing to consider that microsporidia may have lost Rb from their genome while simultaneously inhibiting Rb in their host, converting intestinal cells into a more permissive environment for replication. A similar phenomenon has been observed in leprosy bacteria, where host cell de-differentiation enhances bacterial spread (70). In *C. elegans*, somatic tissues of long-lived mutants exhibit germline gene expression, which enhances resistance to genotoxic stress and improves healthspan (71). This soma-to-germline transformation during infection might boost pathogen replication while also extending host survival despite the parasite load. Reprogramming host cells could be an efficient strategy for microsporidia to maximize resources while maintaining an intracellular niche for an extended period.

Because loss of the *C. elegans* Rb ortholog, *lin-35*, leads to the upregulation of a gene set similar to those induced during microsporidia infection, it raises the possibility that the host recognizes DRM complex dysfunction as a signal associated with pathogen presence, triggering an immune response. This suggests that SynMuv proteins may have been evolutionarily selected as “guardees” frequently targeted by pathogens and are monitored by the immune system as infection signals. In this context, it parallels the Guard Hypothesis from plant and mammalian immunity (72–74), where immune receptors detect disruptions in key host proteins as a sign of infection. For example, the primary antiviral function of the DNA-binding protein MORC3 in mammalian cells is safeguarded by its secondary role in repressing IFN; thus, a virus that degrades MORC3 to evade its antiviral effects will unleash this secondary immune response (13).

A related hypothesis is that pathogen effectors modify chromatin as a strategy to evade the immune response. For example, the microsporidian *Encephalitozoon hellem* secretes EnP1 into the host’s nucleus, inhibiting histone 2B monoubiquitylation and suppressing p53-mediated ferroptosis (75). Furthermore, adenovirus counters host defenses through chromatin remodeling, with its early protein E1A forming a p300-e1a-RB1 complex that condenses chromatin, repressing host genes by redistributing histone modifications (76, 77). These chromatin changes could act as tripwires, triggering immune system activation. In fact, chromatin modifications are tightly linked to immune regulation, as nucleosome remodeling is essential for activating pathogen response genes and IFN-I-stimulated factors (78, 79). Additionally, Rb inhibits IFN-β production by deacetylating its promoter (80), while the NuRD complex downregulates STAT1 through histone deacetylation (81). These insights underscore the critical role of chromatin regulators in fine-tuning immune responses and maintaining proper immune function.

Most of the genes we investigated belong to the SynMuvB class of SynMuv gene family, but our RNAi screen also identified several genes previously classified as SynMuvA (*smo-1* and *uba-2*) and SynMuvC (*mys-1* and *ttr-1*) (47, 61, 82). These three classes work in parallel pathways to repress inappropriate vulval cell fates (47). Our results suggests that multiple SynMuv classes may be involved in immune regulation. Notably, while the SUMOylation genes *smo-1* and *uba-2* are categorized as SynMuvA, they exhibit functional characteristics of both SynMuvA and SynMuvB classes in vulval development (47). Similarly, the SynMuvB gene *ubc-9* also displays SynMuvA-like functions in vulval development (47). The remaining core SUMOylation components, AOS-1 and GEI-17, are not classified as SynMuv genes, yet we found that all SUMOylation genes negatively regulate the IPR. We hypothesize that, like other SynMuv genes, SUMOylation genes contribute to chromatin remodeling during IPR activation. This role might be similar to the function of SUMOylation in mammals, where the SUMOylated chromatin-modifying enzyme SETDB1 (SET domain bifurcated histone lysine methyltransferase 1) mediates H3K9me3 methylation, causing chromatin compaction and repressing IFN-I response activation (83).

SUMOylation plays several crucial roles in development. In addition to its function in vulval formation, SUMO components are essential for normal organismal development, as their downregulation can cause embryonic arrest or significantly delay development (84, 85). Interestingly, mutations in other SynMuv genes also result in developmental abnormalities. Although the DRM complex negatively regulates the cell cycle, mutations in the DRM gene *lin-54* and the associated gene *lin-15B* cause delayed intestinal cell divisions and development (86). In this study, we observed delayed postembryonic development in *lin-15B* mutants. Given that prolonged activation of the IPR and IFN-I responses has been linked to delayed development and developmental arrest (7, 21, 26), it is possible that the delayed development in SynMuv mutants may in part be due to immune system overactivation – an aspect that has not been previously explored. In conclusion, our study identified numerous genes involved in regulating innate immunity in *C. elegans*. Importantly, most of these genes are conserved in mammals. Given the regulatory similarities between the IPR and IFN-I response (7), it is possible that these genes retained their immunosuppressive functions across species.

## MATERIALS AND METHODS

### C. elegans strains

*C. elegans* strains were cultured on Nematode Growth Media (NGM) agar plates seeded with streptomycin-resistant *Escherichia coli* OP50-1 and grown at 20°C, unless otherwise specified (87). The strains utilized in this study are listed in S5 Table.

### Synchronization of *C. elegans*

*C. elegans* population synchronization was performed as described before (88). In brief, gravid adult animals were washed off plates with M9 buffer and collected into 15 ml conical tubes. The tubes were centrifuged, and the supernatant was removed, leaving 3 ml of M9 buffer. 1 ml of bleaching solution (500 μl of 5.65–6% sodium hypochlorite and 500 μl of 5 M NaOH) was added, and the tubes were mixed until the adult animals were partially dissolved. The volume was brought up to 15 ml with M9 buffer to wash the released embryos. The tubes were immediately centrifuged, and the supernatant was removed. This washing step was repeated four times, and the embryos were resuspended in a final volume of 3 ml of M9 buffer. The embryos were then incubated at 20 °C with continuous rotation for 18–24 hours to allow them to hatch into L1 larvae.

### Forward genetic screens and mapping of *lin-15B* alleles

The forward genetic screens and allele mapping were performed as previously described (26, 29, 89). In brief, L4-stage animals carrying the *jyIs8[pals-5p::gfp, myo-2p::mCherry]* transgene (and *pals-22(jy1) pals-25(jy11)* mutations from the first screen, where the *jy76* allele was isolated, and wild-type animals from the second screen, where the *jy86* allele was isolated) were harvested from NGM plates using M9 buffer. The worms were then collected into 15 mL conical tubes, centrifuged, and the supernatant was discarded. Afterward, the worms were washed twice with M9 to remove residual bacteria. The pellet was resuspended in 4 ml of M9, and 25 μl of mutagen, ethyl methanesulfonate (EMS), was added. The worms were incubated at room temperature with continuous rotation for 4 hours. Post-incubation, the worms were centrifuged again, and the supernatant was discarded. They were washed twice more with M9 to remove the mutagen. The treated worms were then plated on 10 cm Petri dishes containing OP50-1 bacteria and incubated at room temperature for 2.5 hours to recover. Following this recovery period, mutagenized worms were transferred to fresh 10 cm Petri plates with OP50-1 bacteria and incubated at 20 °C. After 36 hours, the P0 animals were removed, and the F1 progeny continued to develop. After three days, F1 adults were bleached to synchronize their F2 progeny, which were then screened for *pals-5*p::GFP expression. The *jy86* allele was identified in the F2 population, while the *jy76* allele was identified in the F3 generation following F2 bleaching. Approximately 33,500 and 26,000 haploid genomes were screened in the first and second screens, respectively. Isolated mutants were backcrossed six times to the wild-type strain. Backcrossing to the original strains used in the screens revealed that the causative mutation is located on the X chromosome. In a complementation test, *jy76* and *jy86* failed to complement each other, suggesting they carry causative mutations in the same gene.

Genomic DNA was extracted from both wild-type and mutant strains using the Qiagen Puregene Core Kit A and sent to BGI Genomics for whole genome sequencing. Data analysis was conducted using the Galaxy platform (https://usegalaxy.org). Read quality was evaluated with FastQC. Reads were aligned to the *C. elegans* reference genome (WBCel235.dna.toplevel.fa.gz) using Bowtie2 (90, 91). Genetic variants were detected with FreeBayes. Allelic variants were processed with VcfAllelicPrimitives to split allelic primitives into separate VCF lines. The VCF datasets for wild-type and mutants were compared using VCF-VCFintersect. Variants were filtered based on the criteria (QUAL > 30) & (DP ≥ 10) & (isHOM(Gen[0])) using SnpSift Filter (92), and the remaining variants were annotated with the SnpEff eff program (93).

### Imaging

For the analysis of *pals-5*p::GFP expression, animals were anesthetized with either 20 or 100 µM levamisole. They were then placed on agarose pads on glass slides and covered with cover slips. The samples were imaged using a Zeiss Axio Imager.M2 compound microscope with ZEN 3.8 software or a Zeiss Axio Imager.M1 compound microscope with Axio Vision 4.8.2 software. Image processing was performed using the FIJI software (94).

### Single-molecule fluorescence in situ hybridization (smFISH)

Synchronized animals were placed on NGM plates seeded with OP50-1 bacteria and incubated at 20°C for 48 hours. After incubation, the animals were washed off the plates and fixed in 4% paraformaldehyde with PBST (phosphate-buffered saline containing 0.1% Tween 20) for 30 minutes at room temperature. Following fixation, the samples were incubated overnight at 4°C in 70% ethanol. For staining, 1 µM Quasar 670-conjugated *pals-5* smFISH probes (Biosearch Technologies) were used in a hybridization buffer containing 10% formamide, 2× SSC, 10% dextran sulfate, 2 mM vanadyl ribonucleoside complex, 0.02% RNase-free BSA, and 50 µg of *E. coli* tRNA. The samples were incubated overnight at 30°C in the dark. Afterward, they were washed with a buffer containing 10% formamide and 2x SSC at 30°C in the dark for 30 minutes. Vectashield with DAPI was then added to each sample, and the stained animals were transferred onto microscope slides, covered with glass coverslips, and imaged. Z-stack images of the region containing the anterior part of the intestine were acquired using a Zeiss Axio Imager.M1 compound microscope equipped with a 63x oil immersion objective and Axio Vision 4.8.2 software. Z stack processing into maximal projection images was performed using the FIJI software (94).

### Microsporidia infection

One million *N. parisii* spores were combined with 10 µl of 10x OP50-1 *E. coli*, along with 1,200 synchronized L1-stage *C. elegans*, and M9 buffer, bringing the total volume to 300 µl. This mixture was spread on 6 cm NGM plates, left to dry for 30 minutes at room temperature, and then incubated at 25 °C for 3 hours. Following incubation, the worms were collected using M9 buffer with 0.1% Tween 20, washed in PBST, and fixed in 4% paraformaldehyde for 45 minutes. The fixed worms were stained overnight at 46°C using Cal Fluor 610 red FISH probes labeled with the fluorophore, that specifically bind to *N. parisii* ribosomal RNA (95). Imaging was performed with a Zeiss Axio Imager.M1 compound microscope with Axio Vision 4.8.2 software, and data were analyzed and statistically processed using GraphPad Prism 10 software.

### Orsay virus infection and quantification of viral RNA1

Approximately 2,000 synchronized L1-stage worms were placed on 10 cm NGM plates seeded with *E. coli* OP50-1 for each strain. The plates were kept at 20°C for about 44 hours until the worms reached the L4 stage. A viral mixture was prepared by diluting 30 μl of the Orsay virus filtrate (1:10 in M9 buffer) and combining it with 150 μl of concentrated OP50-1 and 600 μl of M9 buffer. This mixture was added to 10 cm plates containing L4-stage worms, dried at room temperature, and incubated at 20°C for 24 hours. RNA was extracted using Tri-reagent (Molecular Research Center, Inc) and BCP phase separation reagent (Molecular Research Center, Inc), and converted to cDNA with the iScript cDNA synthesis kit (Bio-Rad). For qRT-PCR, iQ SYBR Green Supermix (Bio-Rad) was used along with primers specific to the Orsay virus RNA1 (S6 Table). Gene expression was normalized to *snb-1*, which remained constant under the tested conditions. Each sample was run in two technical replicates, and three independent experimental replicates were performed.

### Bead-feeding assay

The bead feeding assay was performed as previously described (26). Briefly, 1,200 synchronized L1-stage worms were combined with 6 µl of fluorescent beads (Fluoresbrite Polychromatic Red Microspheres, Polysciences Inc.), 25 µl of a 10X concentrated OP50-1 *E. coli* solution, and M9 buffer to a total volume of 300 µl. This mixture was spread onto 6 cm NGM plates, allowed to air dry for 5 minutes, and incubated at 25°C. After a 5-minute incubation, plates were placed on ice. Worms were washed off the plates with ice-cold PBST, then fixed in 4% paraformaldehyde for 30 minutes. Following fixation, the samples were washed with PBST, and the worms were imaged using a Zeiss Axio Imager.M2 compound microscope with ZEN 3.8 software. Red fluorescence was quantified in the FIJI program (94), analyzing 50 worms per strain in each of three experimental replicates. Data were processed with GraphPad Prism 10.

### Transcriptomics and ChIP-seq data analyses

Data were obtained from the publications referenced in the main text. Data overlap was analyzed using Multiple List Comparator software (https://molbiotools.com/listcompare.php). The gene name conversions were made using WormBase ParaSite Biomart (https://parasite.wormbase.org/biomart).

### RNA interference screen

RNA interference (RNAi) assays were conducted using the feeding method (96). 23 distinct overnight cultures of HT115 *E. coli* RNAi strains were seeded onto RNAi plates (NGM plates supplemented with 5 mM IPTG and 1 mM carbenicillin) and incubated at room temperature for 3 days. L4440 served as a negative RNAi control, while *pals-17* RNAi was used as a positive control for inducing *pals-5*p::GFP expression. L4-stage *C. elegans* were transferred to RNAi plates and incubated at 20°C. After 48 and 72 hours, *pals-5*p::GFP expression was evaluated in the F1 progeny, with separate analysis for L1-L3 stage animals and L4-young adult stage animals. A total of 100 animals were analyzed per age group and per RNAi condition. Phenotypic analysis was performed using a Stereo Discovery V8 fluorescent microscope equipped with an X-Cite XYLIS XT720S illumination system. Data analysis was carried out using GraphPad Prism 10 software.

### SUMO RNAi treatments and qRT-PCR analysis of IPR gene expression

RNA interference assays were performed using the feeding method (96). Overnight cultures of *E. coli* were plated on RNAi plates (NGM plates supplemented with 5 mM IPTG and 1 mM carbenicillin) and incubated at room temperature for 24 hours. L4 animals were picked onto 3 10-cm seeded RNAi plates per strain (number of animals per plate: L4440 control vector RNAi – 12; *aos-1* RNAi – 20; *gei-17* – 18, *ubc-9* – 15; *smo-1* – 24) and incubated at 20 °C until majority of F1 progeny reached the L4 stage. RNA isolation was performed as previously described using Tri-reagent (Molecular Research Center, Inc) and BCP phase separation reagent (Molecular Research Center, Inc), and cDNA synthesis was performed using the iScript DNA synthesis kit (Bio-Rad) (26). qRT-PCR was performed using iQ SYBR Green Supermix (Bio-Rad) with the CFX Connect Real-Time PCR Detection System (Bio-Rad). As a control, values were normalized to expression levels of the neuronal gene *snb-1*, which is not affected by IPR activation. Primer sequences are listed in S6 Table. Data analysis was performed through PRISM using the Pfaffl method. One-tailed t-tests were used to determine statistical significance between samples. Each sample was run in two technical replicates, and three independent experimental replicates were performed.

### Yeast two-hybrid analysis

The yeast two-hybrid (Y2H) screen was carried out using the ULTImate Y2H cell-to-cell mating method (Hybrigenics Service, Evry, France), as previously described (27). In brief, the coding sequence for PALS-20 was synthesized and inserted into the pB66 plasmid downstream of the Gal4 DNA-binding domain. This construct served as the bait to screen a *C. elegans* mixed-stage cDNA prey library, which was generated through random priming and cloned into the pP6 plasmid containing the Gal4 activation domain. The screen employed a haploid mating strategy, with one yeast sex type (mata) containing the bait and the other (matα) carrying the prey constructs (97). In total, 79.8 million interactions were examined, yielding 279 clones that grew on selective media lacking tryptophan, leucine, and histidine. Prey fragments from the positive clones were PCR-amplified and sequenced at both 5’ and 3’ ends to identify the corresponding proteins using the NCBI GenBank database. The Predicted Biological Scores (PBS), which assess the reliability of bait-prey interactions, were explained previously (27, 98) and are summarized for all identified interactions in S5 Table.

## ACKNOWLEDGEMENTS

This work was funded by NIH grants R01 AG052622, GM114139, and AI176639 and NSF grant 2301657 awarded to E.R.T., the American Heart Association postdoctoral award 19POST34460023 and The George Washington University Facilitating Fund Award to V.L., and NIGMS/NIH Institutional Research and Academic Career Development Award K12GM068524 to E.T. We appreciate the helpful manuscript feedback from James Tirtorahardjo and Lakshmi Batachari. The model figures were created using BioRender.com.

---

**S1 Fig.**
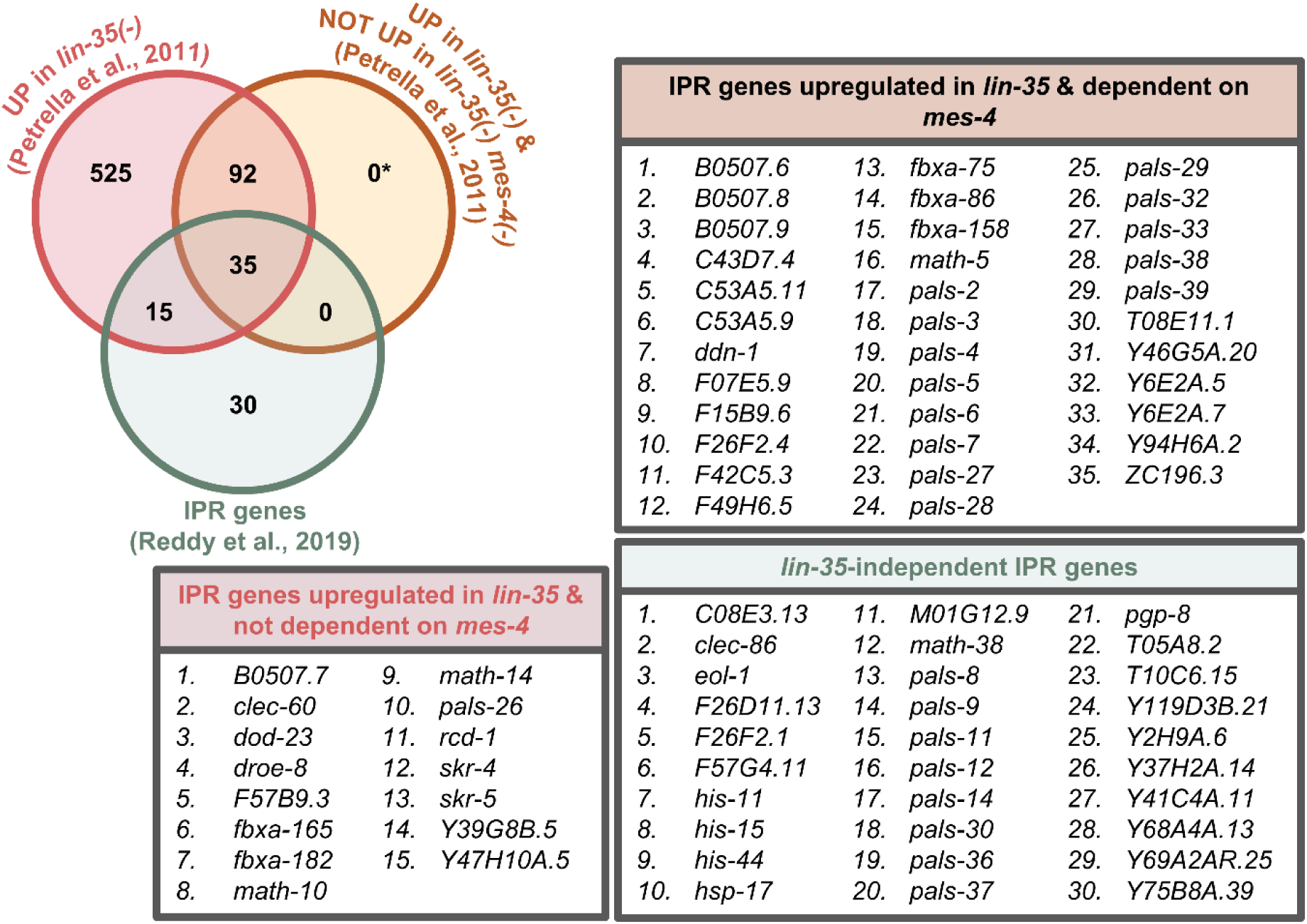
Most of IPR genes upregulated in *lin-35* mutants are dependent on *mes-4*. *In silico* analysis of previously published transcriptomic data. The asterisk indicates a gene that was removed from the original list because it was not found among the genes upregulated in the lin-35(-) mutant.

**S2 Fig.**
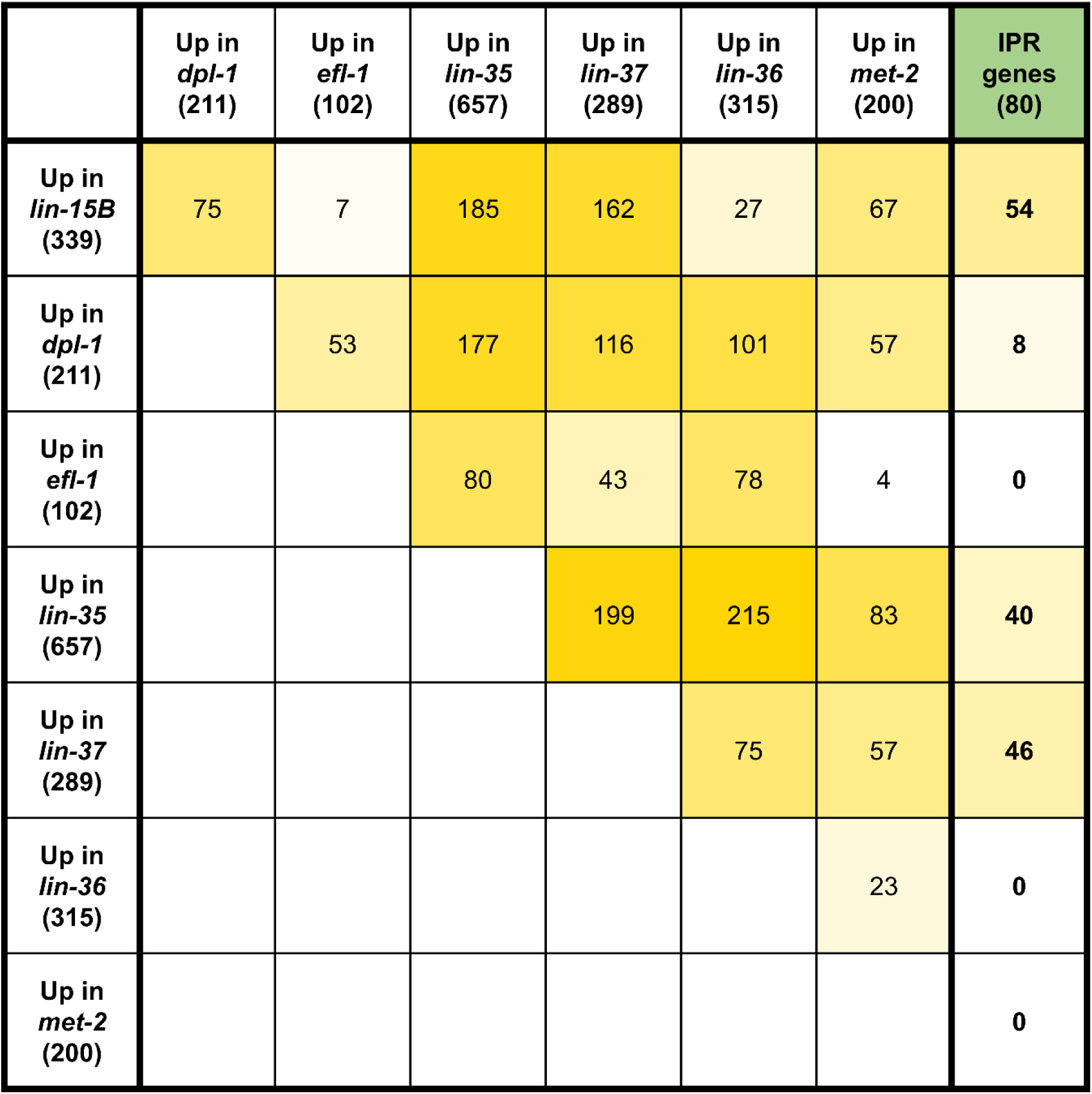
A subset of SynMuv genes regulate IPR gene expression. *In silico* analysis a previously published transcriptomic datasets. The numbers in parentheses indicate the count of upregulated genes in the annotated genetic backgrounds and IPR genes. Numbers in the intersecting cells represent the number of genes common to both datasets.

**S1 Table. Comparative analysis of IPR genes and transcriptomic data for *lin-15*, *lin-35* and *mes-4*.**

**S2 Table. Comparative analysis of IPR genes and transcriptomic and ChIP-seq data for *lin-15B*/LIN-15B.**

**S3 Table. Comparative analysis of IPR genes and transcriptomic data for several SynMuv mutants.**

**S4 Table. Yeast two-hybrid analysis of PALS-20 and PALS-25.**

**S5 Table. List of strains used in this study.**

**S6 Table. List of primers used in this study.**

